# MRE11-RAD50-NBS1 activates Fanconi Anemia R-loop suppression at transcription-replication conflicts

**DOI:** 10.1101/472654

**Authors:** Emily Yun-chia Chang, James P. Wells, Shu-Huei Tsai, Yan Coulombe, Yujia A. Chan, Yi Dan Zhu, Louis-Alexandre Fournier, Philip Hieter, Jean-Yves Masson, Peter C. Stirling

## Abstract

Ectopic R-loop accumulation causes DNA replication stress and genome instability. To avoid these outcomes, cells possess a range of anti-R-loop mechanisms, including RNaseH that degrades the RNA moiety in R-loops. To comprehensively identify anti-R-loop mechanisms, we performed a genome-wide trigenic interaction screen in yeast lacking *RNH1* and *RNH201*. We identified >100 genes critical for fitness in the absence of RNaseH, which were enriched for DNA replication fork maintenance factors such as *RAD50*. We show in yeast and human cells that R-loops accumulate during RAD50 depletion. In human cancer cell models, we find that RAD50 and its partners in the MRE11-RAD50-NBS1 complex regulate R-loop-associated DNA damage and replication stress. We show that a non-nucleolytic function of MRE11 is important for R-loop suppression via activation of PCNA-ubiquitination by RAD18 and recruiting anti-R-loop helicases in the Fanconi Anemia pathway. This work establishes a novel role for MRE11-RAD50-NBS1 in directing tolerance mechanisms of transcription-replication conflicts.

## Introduction

Genome instability refers to a state in which cells experience a higher than normal mutational burden during division. Importantly, genome instability is an early driver of tumourigenesis because it increases the chance that oncogenic mutations will occur ^1^. Genome instability can arise from increased DNA damage, reduced DNA replication or mitotic fidelity, or perturbed cell cycle checkpoints. The interplay of these cellular pathways, their interactions with environmental insults, and their role in cancer formation, evolution, and treatment are an active area of inquiry.

One genome instability mechanism of growing importance is genotoxic transcription-replication collisions ^2^. Such interference happens normally, and cells have evolved genome architectures and replication fork stabilizing proteins that help to mitigate the risks of transcription to the genome. Nonetheless, when transcription is aberrant, or the replication fork is not robust to stalling, collisions can lead to DNA breaks, which, in excess, can promote error prone repair mechanisms and mutations. A special case of this phenomenon involves extended annealing of nascent RNA to the template DNA strand to create a three-stranded nucleic acid structure called an R-loop. R-loops perform important functions in transcriptional regulation, but can also act as barriers to replication fork progression when they accumulate ectopically through a mechanism that is not completely understood^3^.

In order to prevent R-loop accumulation and the resulting DNA replication stress, cells have evolved a host of R-loop degrading and mitigating strategies. For instance, splicing, RNA processing, and RNA packaging machinery sequester nascent RNAs to prevent them from reannealing to the genome ^4–6^; topological regulators such as Topoisomerase I and II restrict access of genomic DNA to RNA transcripts^7,8^; RNA nucleases including RNaseH1 and RNaseH2 specifically target the RNA moiety of R-loops^9^; Senataxin, Aquarius and other helicases have been implicated in DNA:RNA hybrid unwinding; and DNA repair proteins such as XPF and XPG have been suggested to directly cleave R-loops to promote their resolution^10,11^. More recently, core components of the replication fork and associated factors have been linked to R-loop resolution, although the mechanisms at play are currently unclear. Notably, these include several proteins related to the Fanconi Anemia pathway, some of which have helicase activity which could remove R-loops ^12–14^.

The MRE11-RAD50-NBS1 (MRN) complex is a highly conserved genome maintenance protein complex. MRN plays catalytic roles in genome stability through the nuclease activity of MRE11, as well as structural roles where the complex can serve as both a DNA tether and a platform for DNA damage signaling^15^. MRN binds to DNA ends and regulates early steps of the double-strand break repair pathway, and plays an important role in DNA end resection in collaboration with other nucleases ^15^. In addition, MRN has at least two important roles at stalled DNA replication forks. In BRCA2-deficient genetic backgrounds, MRE11 cleaves nascent DNA and promotes fork degradation-based restart mechanisms^16,17^. The related yeast MRX (Mre11-Rad50-Xrrs2) complex functions in a nuclease-independent manner to tether sister chromatids together at stalled forks and promote efficient fork restart through an interaction with a single-stranded DNA binding protein^18,19^.

In an effort to identify new players in cellular R-loop tolerance, we conducted a screen for genes that become essential for robust growth in yeast lacking RNaseH enzymes, which are encoded by *RNH1* and *RNH201*. We find a strong enrichment of DNA replication fork-related processes, including members of the MRX/MRN complex. In human cell models, we find that MRN functions to suppress R-loop-associated DNA damage and DNA replication stress in a manner that is independent of its nucleolytic activity. Instead, MRN is important for the recruitment of Fanconi Anemia pathway proteins to R-loop sites by potentially impacting PCNA-ubiquitination levels. Together these data define a new role for MRN in coordinating the response to an endogenous source of genome instability during DNA replication.

## Results

### Identification of RNaseH-dependent yeast mutants

RNaseH1, encoded by *RNH1*, and RNaseH2, a trimeric complex with a catalytic subunit encoded by *RNH201*, exert semi-redundant functions in genome maintenance^20^. While both enzymes can digest the RNA moiety in R-loops, Rnh201 also functions in ribonucleotide-excision repair (RER) and Okazaki fragment processing, while Rnh1 has mitochondrial functions^9,21^. We reasoned that mutants which specifically require the shared R-loop resolving activity will be less fit as triple mutants (*genex*Δ*rnh1*Δ*rnh201*Δ) in comparison with either double mutant combination (*genex*Δ*rnh1*Δ or *genex*Δ*rnh201*Δ). To conduct this experiment, we used synthetic genetic array (SGA^22^) with an *rnh1*Δ*rnh201*Δ double mutant query strain (rnhΔΔ), and generated genetic interaction profiles for each double mutant and triple mutant combination (**Table S1**). Our screen revealed fewer significant interactions for *rnh1*Δ (23) than for *rnh201*Δ (37), and a much larger group for *rnh*ΔΔ (119), conveying the strong buffering relationship between Rnh1 and Rnh201 (**Fig. 1A**). While a majority of *rnh201Δ* interactions were shared with *rnh*ΔΔ, there were 95 candidate negative-genetic interactions unique to *rnh*ΔΔ (**Fig. 1**). Gene ontology (GO) analysis of the raw *rnh*ΔΔ negative genetic interaction list revealed a set of highly enriched terms entirely directed at DNA repair and DNA replication (**Fig. 1B**). Highly enriched cellular components include the *Mre11 complex, Smc5/6 complex*, and *Replication Fork*, while enriched Biological Processes include *DNA strand elongation* and *intra-S DNA damage checkpoint*. These suggest that losing R-loop mitigating activity in the *rnh*ΔΔ strain places a significant burden on normal replication, which is evident from increased fitness defects when replication is further perturbed. These findings cohere with recent reports characterizing the *rnh*ΔΔ strain in the presence of conditional Topoisomerase I inactivation^23^, and with the known role of R-loops in causing replication stress^3,7^.

**Figure 1.**
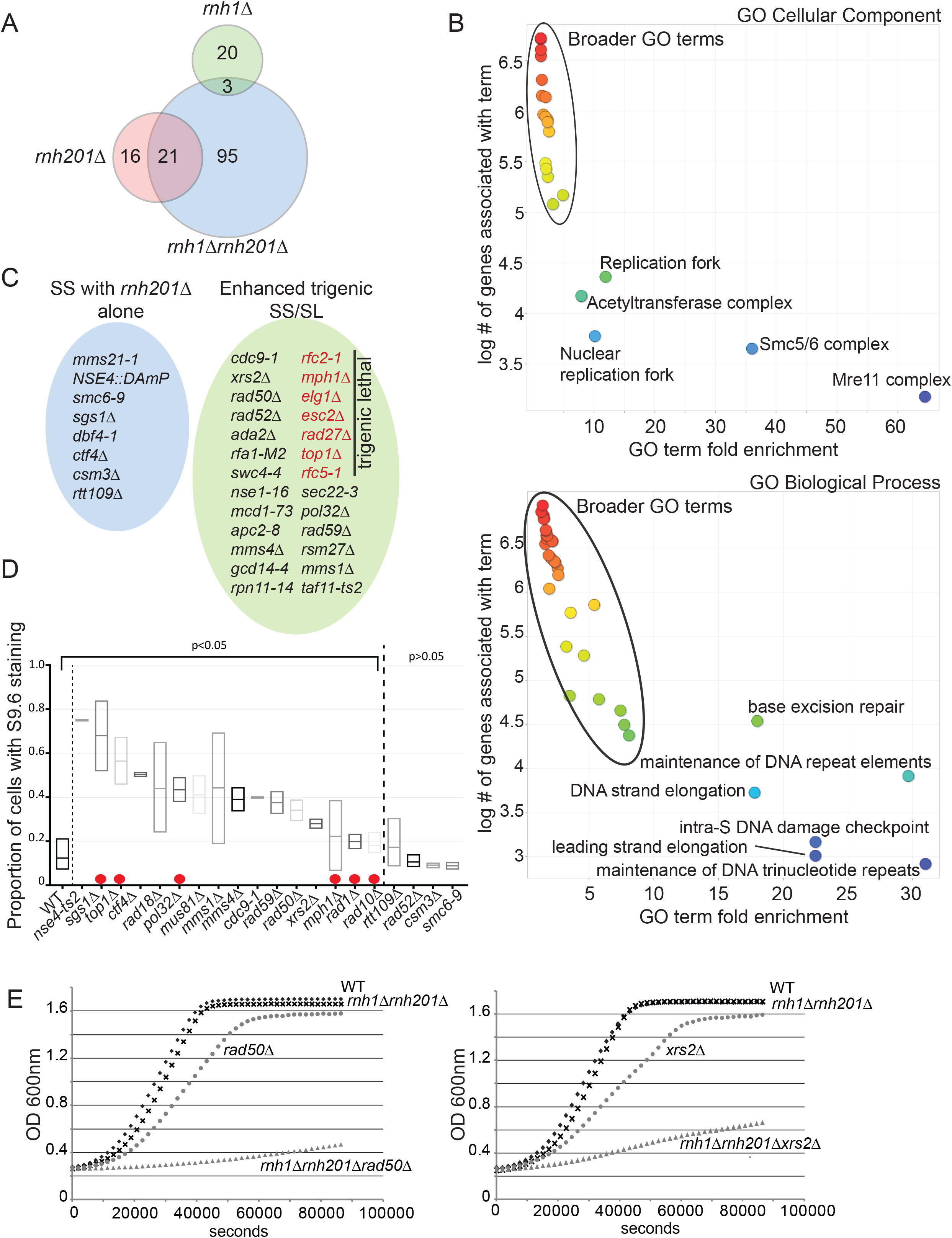
RNaseH-deficient yeast depend on replisome function and fork protection for fitness. (A) Negative genetic interaction results of double and triple mutant SGA screens with the indicated query genotype. A Venn diagram of candidate negative genetic interactions is shown. (B) Gene Ontology analysis of negative *rnh*ΔΔ interacting partners. Plots show the output of ReviGO, which effectively trims redundant GO terms. (C) Validated interaction partners by tetrad analysis and growth curves. Synthetic sick (SS) interactions with *rnh201*Δ are shown in blue circle, genes that exhibited enhanced sickness or lethality (SL) in *rnh*ΔΔ are shown in green circle. Gene deletions in red were lethal in the context of the *rnh*ΔΔ genotype. (D) S9.6 staining of chromosome spreads in the indicated strains. Strains between the dotted lines had a significantly higher proportion of stained nuclei than WT (Fisher Exact Test. p<0.05, Holm-Bonferroni corrected). Red dots indicate predicted positive controls (see main text). (E) Representative growth curves of *rad50Δ* or *xrs2*Δ with *rnh*ΔΔ. Statistical analysis of area under the curve for triplicates confirm that the triple mutants are significantly sicker than expected, p<0.001.

We validated the genetic interactions for a selection of hits representing the enriched GO terms, and found that while some mutants were already synthetic sick (SS) when *RNH201* was absent, many were considerably sicker or inviable when *RNH1* was also absent (**Fig. 1C**). We then directly tested the accumulation of R-loops in twenty candidate R-loop regulatory mutant strains by staining chromosome spreads with the S9.6 antibody that targets DNA:RNA hybrids (**Fig. 1D**)^24,25^. This small screen included six known or predicted positive controls (red dots in **Fig. 1D**) based on studies in yeast (SGS1, TOP1, MPH^18,14,26^), or of human orthologues (RAD1/10, POL32)^10,27^. Gratifyingly, all six strains showed significant increases in S9.6 staining providing additional confidence in the assay. Collectively, this experiment determined that some mutations in the SMC5/6 complex (*NSE4*) and multiple DNA replication-associated repair factors (*SGS1, TOP1, RAD18, POL32, MUS81/MMS4, RAD50, XRS2, & MPH1*) were both negative genetic interactors of *rnhΔΔ* and also accumulated DNA:RNA hybrids (**Fig. 1D**). Such observations has been previously reported for deletions of *SGS1, TOP1*, and *MPH1*^7,12,14^ but a role for the Mre11-Rad50-Xrs2 complex in R-loop tolerance has not been described. Quantitative growth curve analysis of triple mutants of *rnh*ΔΔ and either *rad50*Δ and *xrs2*Δ illustrated the dramatic loss of fitness associated with dual MRX and RNaseH loss (**Fig. 1E**). Given the potential role of MRN as a tumour-suppressor complex in breast and other cancers ^28^, we wondered whether this phenotype was conserved to human cancer cell models.

### R-loop accumulation and RNaseH-dependent DNA damage in MRN complex depleted cancer cell lines

We first tested whether human RNaseH2A knockout cells exhibited enhanced sickness when depleted for MRN components. siRNA knockdown of MRE11, RAD50 or NBS1 in isogenic RNaseH2A+ or RNaseH2A-HeLa cells^29^ showed selective toxicity in the knockout cells, consistent with a conserved genetic interaction between MRN and RNaseH (**Fig. 2A and Fig. S1A/S1B**). We next tested whether MRN depletion increased R-loop accumulation. S9.6 staining in cells with MRN subunit depletion showed significant increases in R-loop accumulation (**Fig. 2B and Fig. S1C**), and this was suppressed by cotransfection with RNaseH1-GFP (**Fig. 2C**) ^30^. Indeed, DNA:RNA immunoprecipitation (DRIP) experiments revealed an induction of hybrid occupancy at R-loop-prone loci ^31^ when RAD50 was depleted (**Fig. 2D**).

**Figure 2.**
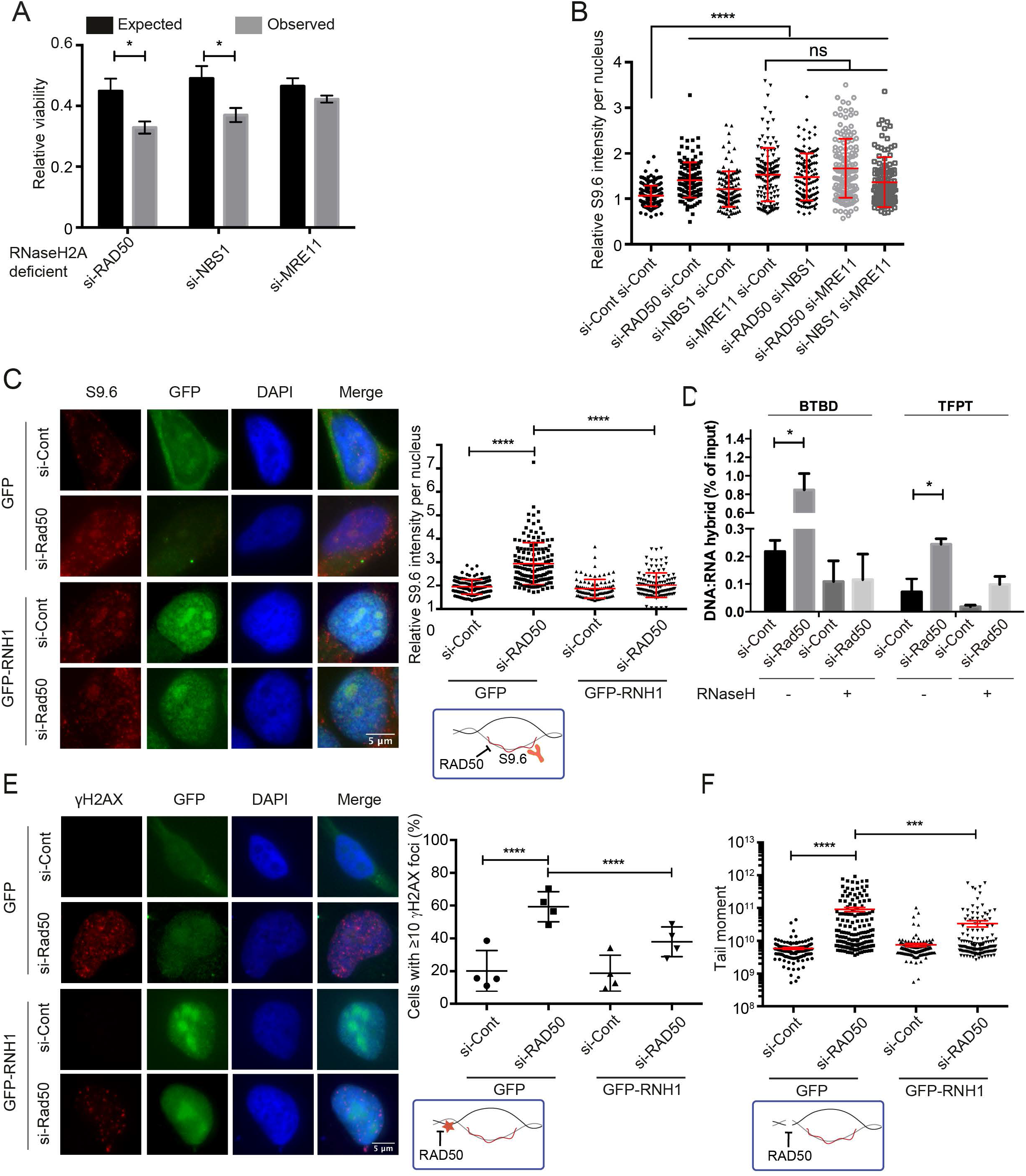
Conserved RNaseH-dependence and R-loop accumulation in human cells depleted for the MRN complex. (A) Cell viability of RNaseH2A normal and knockout cells depleted for *RAD50, MRE11* or *NBS1* as measured by crystal violet stain (N = 4; *t* test; mean ± SEM) showed observed viability values were significantly lower than expected values after depletion of *RAD50* and *NBS1*. (B) Relative S9.6 staining intensity per nucleus in MRN complex depleted cells. N = 3; ANOVA; mean ± SD. (C) Representative images (left) and quantification (right) of S9.6 staining per nucleus in HeLa cells treated with si-Cont or si-RAD50. Cells were transfected with either a control vector (GFP) or one expressing GFP-RNaseH1 (GFP-RNH1). N = 3; *t* test; mean ± SD. (D) DRIP-qPCR analysis of R-loop accumulation at BTBD and TFPT loci in control and si-RAD50 cells with or without RNaseH treatment before precipitation. N = 3; *t* test; mean ± SEM. (E and F) RNaseH1-dependent DNA damage phenotypes in RAD50-depleted cells shown by γ-H2AX staining (E) (N = 4; Fisher’s exact test; mean ± SD) and neutral comet assay for DNA breaks (F) (N = 3; *t* test; mean ± SEM). *, P < 0.05; ****, P < 0.0001. For C-F and some panels in subsequent figures (3A,B, 5 and 6) cartoon schematics of an R-loop illustrate the effects being tested by the experiment.

To determine whether the R-loop accumulation we observed contributed to DNA damage phenotypes we knocked down RAD50, which ablates complex function, and tested the R-loop-dependence of DNA damage phenotypes. Significant DNA damage induction associated with RAD50 depletion was observed by quantifying DNA damage signaling by γ-H2AX staining, or measuring DNA double-strand breaks with a neutral comet assay. Importantly, both DNA damage phenotypes were partially suppressed by the overexpression of RNaseH1 (**Fig. 2E, Fig. 2F and Fig S1D**). Together, these data show that human cells lacking MRN function depend on RNaseH2A for fitness, and that MRN-depleted cells accumulate R-loops and R-loop-dependent DNA damage. MRN plays both structural and nucleolytic roles in double-strand break repair and replication fork stability maintenance^15^, this raises the question of which pathway impacted by MRN is important for R-loop tolerance and bypass.

### R-loops drive replisome slowing and replication stress in RAD50 depleted cells

One of the best-characterized causes of R-loop-dependent DNA damage is the interference of R-loops with DNA replication fork progression, causing replication stress ^3^. To test the influence of R-loops on these phenotypes in MRN-depleted cells, we first tested the influence of ectopic RNaseH1 expression on the induction of DNA damage signaling in RAD50-depleted cells. Depleting RAD50 caused activation of both the ATM and ATR pathways as judged by Chk2 and Chk1 phosphorylation, respectively (**Fig. 3A**). This also caused phosphorylation of RPA2-ser33, a known ATR-dependent mark of DNA replication stress. As previously reported by other groups, we also saw that ectopic RNaseH1 expression caused activation of DNA damage responses on its own (**Fig. 3A,B**)^32,33^. We believe that this is consistent with RNaseH1 mistakenly targeting normal functions of R-loops. However, RNaseH1-expression reduced Chk1 and RPA2 phosphorylation during RAD50 knockdown, indicating that the ATR-sensitive DNA replication stress response may be R-loop-dependent (**Fig. 3A**). This is consistent with recent literature highlighting the importance of ATR signaling in mediating tolerance to replisome-R-loop conflicts^34,35^. To further explore R-loop-dependent replication stress in RAD50-depleted cells, we conducted native BrdU immunofluorescence to directly assess the exposure of single-stranded DNA during replication stress. RAD50-depleted cells exposed significantly more ssDNA, which was partly suppressed by RNaseH1 (**Fig. 3B**). To more directly measure the frequency of transcription-replication conflicts in MRN-depleted cells, we used a proximity ligation assay (PLA) with antibodies targeting PCNA and RNA polymerase II ^34^: RAD50 depletion significantly increased transcription-replication conflicts marked by PCNA-POLII PLA foci, which again could be reduced by ectopic RNaseH1 expression (**Fig. 3C**). Finally, we conducted DNA combing experiments to directly measure replisome dynamics. As expected, RAD50 depletion led to a net slowing of replication progress, which was partially rescued by transcription inhibition with cordycepin, indicating that transcription impairs replication in RAD50-depleted cells (**Fig. 3D**). To link replisome slowing to R-loops, we overexpressed RNaseH1 which also partially restored replication speed in RAD50-depleted cells (**Fig. 3E**). Overall, our data support a role for the MRN complex in mitigating transcription and R-loop-associated DNA replication slowing and replication stress.

**Figure 3.**
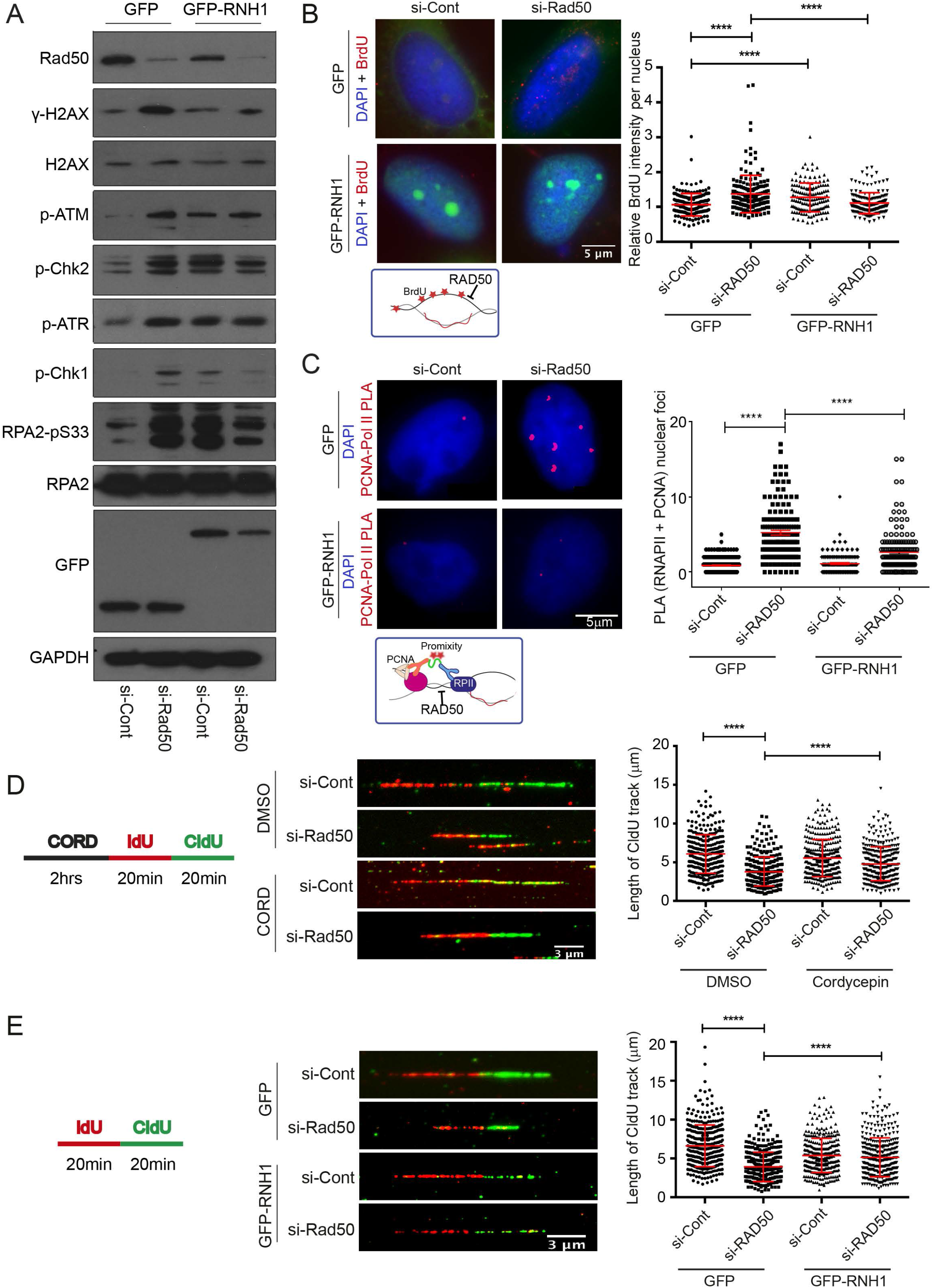
R-loop dependent replication stress in RAD50-depleted cells. (A) DNA damage signaling activation. Representative western blots against the indicated epitope are shown. (B) Native BrdU staining for R-loop dependent ssDNA exposure. Representative images (left) and nuclear fluorescence quantification are shown (right). N = 4; *t* test; mean ± SD. (C) Regulation of transcription-replication conflicts by RAD50. Proximity ligation assay targeting the replisome (anti-PCNA) and RNA polymerase II (anti-RNA Pol II) is shown with representative images (left) and quantification (right), N =3; t-test; mean ± SEM. A cartoon of the assay is below. (D) Transcription dependent replisome slowing in RAD50-depleted cells. Cells were treated with 50 μM cordycepin (CORD) for 2 hours before IdU labeling. (E) R-loop dependent replisome slowing in RAD50-depleted cells. For D and E, an experimental scheme (left), representative DNA fibers from the indicated conditions (middle), and quantified CldU track lengths are shown (right). N = 3; *t* test; mean ± SD. ****, P < 0.0001.

### A non-nucleolytic role for MRN in R-loop regulation

MRN has both catalytic and structural roles in promoting genome stability during DNA replication^17,19,36^. Moreover, through its interactions and roles at replication forks, MRN collaborates with many proteins potentially implicated in R-loop tolerance. For instance, studies describe MRN as recruiting helicases such as BLM^37^, binding WRN^38^, or activating the Fanconi Anemia (FA) pathway ^39^; all of which have been implicated in R-loop tolerance^12–14,40^. To determine whether the catalytic function of MRE11 is important to R-loop tolerance, we treated cells with an MRE11 inhibitor, Mirin ^41^, and found no effect on S9.6 staining in cells (**Fig. 4A**). This suggested that a structural role, rather than a catalytic role for MRN could be important. To test this genetically, we expressed either WT MRE11 or a nuclease dead *mre11^H129N^* mutant in isogenic lymphoblastoid TK6 cell lines, and analyzed S9.6 intensity three days after inducing MRE11 knockdown using 4-hydroxytamoxifen (4-OHT) treatment (**Fig. 4B** and **Fig. S2A**)^42^. This analysis showed that MRE11^−/WT^ or MRE11^−/H129N^ TK6 cells did not show significant differences in S9.6 intensity compared to MRE11^WT/WT^ TK6 cells after 4-OHT treatment or to TK6 cell lines treated with an ethanol control or with no drug treatment. However, MRE11^−/−^ TK6 cells showed a 2-fold increase in mean S9.6 intensity (**Fig. 4B**). These observations suggest that a non-nucleolytic function of MRE11 is important for R-loop regulation. To gain more insights into this, we monitored the activity of the MRN nuclease on DNA/DNA versus RNA/DNA hybrids (**Fig. 4C**). The MRN nuclease activity was very efficient on the DNA/DNA substrates, as nearly 50% of the DNA was degraded at 2.5 nM MRN. In contrast, DNA resection was almost undetectable on RNA/DNA hybrid, leading to a 50-fold decrease in products. Higher concentration of MRN was required to obtain limited degradation of RNA/DNA hybrids (**Fig. 4C**). These studies suggest that MRN nuclease activity is poorly efficient on RNA/DNA, supporting the idea that this activity does not play a major role in R-loop resolution.

**Figure 4.**
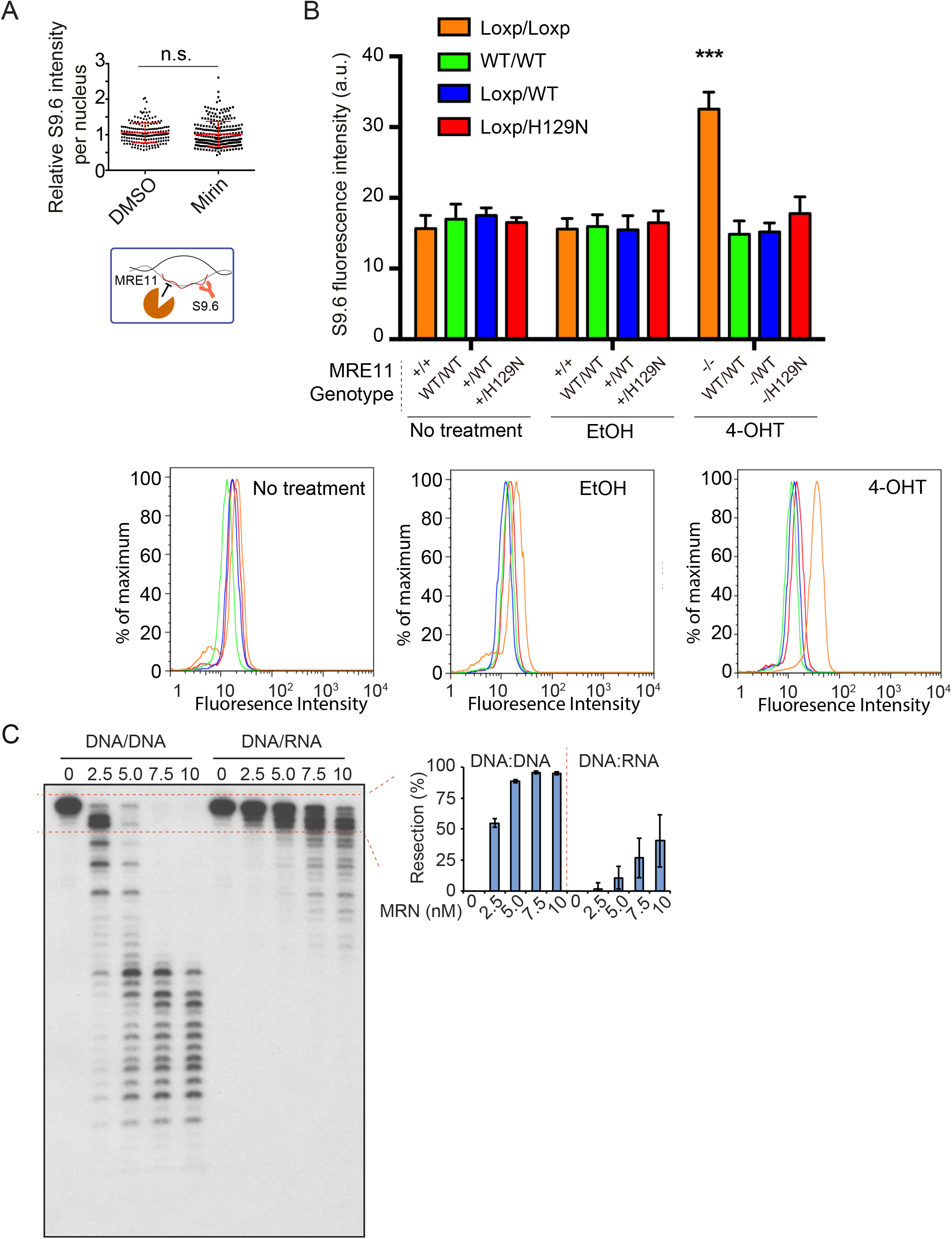
A structural role for the MRN complex in mitigating R-loop associated DNA damage. (A) Nuclear S9.6 staining intensity after treatment with MRE11 inhibitor Mirin (50 μM) is shown. N = 3; *t* test; mean ± SD. A schematic showing the goal of testing MRE11 nuclease activity on R-loops is shown, right. (B) Mean S9.6 fluorescence intensity of TK6 derived lymphoblasts of the indicated genotype after treatment with EtOH (control) or 4-OHT, to induce gene knockout. Representative FACS histograms are shown below, curve colors correspond to bar graph. N = 3; ANOVA; mean ± SEM. ***, P < 0.001. (C) Head-to-head comparison of dsDNA and DNA:RNA hybrid resection by purified MRN complex on 60-mer duplexes. The concentration dependent degradation of the input molecules is plotted on the right.

### MRN functions in the Fanconi Anemia pathway of R-loop resolution

The lack of a direct catalytic role suggested that MRN might function to suppress R-loops through recruitment of other anti-R-loop activities, such as the FA pathway. To determine whether MRN interacts with the FA pathway, we performed an epistasis experiment examining S9.6 staining in cells doubly depleted by siRNA (**Fig. 5A and 5B, Fig. S2B and S2C**). Depletion of MRN, FANCD2, or FANCM gave the expected increase in S9.6 staining, but co-depletion of any of these factors did not further enhance S9.6 staining, suggesting that they work within the same pathway. To confirm that the assay was sufficiently sensitive to observe synergy between separate anti-R-loop pathways, we co-depleted the splicing helicase Aquarius and FANCD2, and observed an additive increase in R-loop levels (**Fig. 5A**). This supports previous observations linking MRN to FA pathway activation^39^, but places these effects in the context of R-loop tolerance for the first time. To further determine whether MRN functions upstream of FA pathway activation we measured the frequency of FANCD2 nuclear foci in cells depleted for RAD50, or as a control, RAD18 which has previously been implicated in FA activation^43^. Remarkably, FANCD2 foci numbers were significantly reduced in RAD50 depleted cells relative to controls (**Fig. 5C**). Interestingly, co-depletion of RAD50 and RAD18 did not significantly further reduce FANCD2 foci (**Fig. 5C** and see below).Thus, despite perturbed replication and excess DNA damage in RAD50-depleted cells (**Fig. 2-3**), FANCD2 recruitment to foci is impaired suggesting a role for MRN upstream of FA.

**Figure 5.**
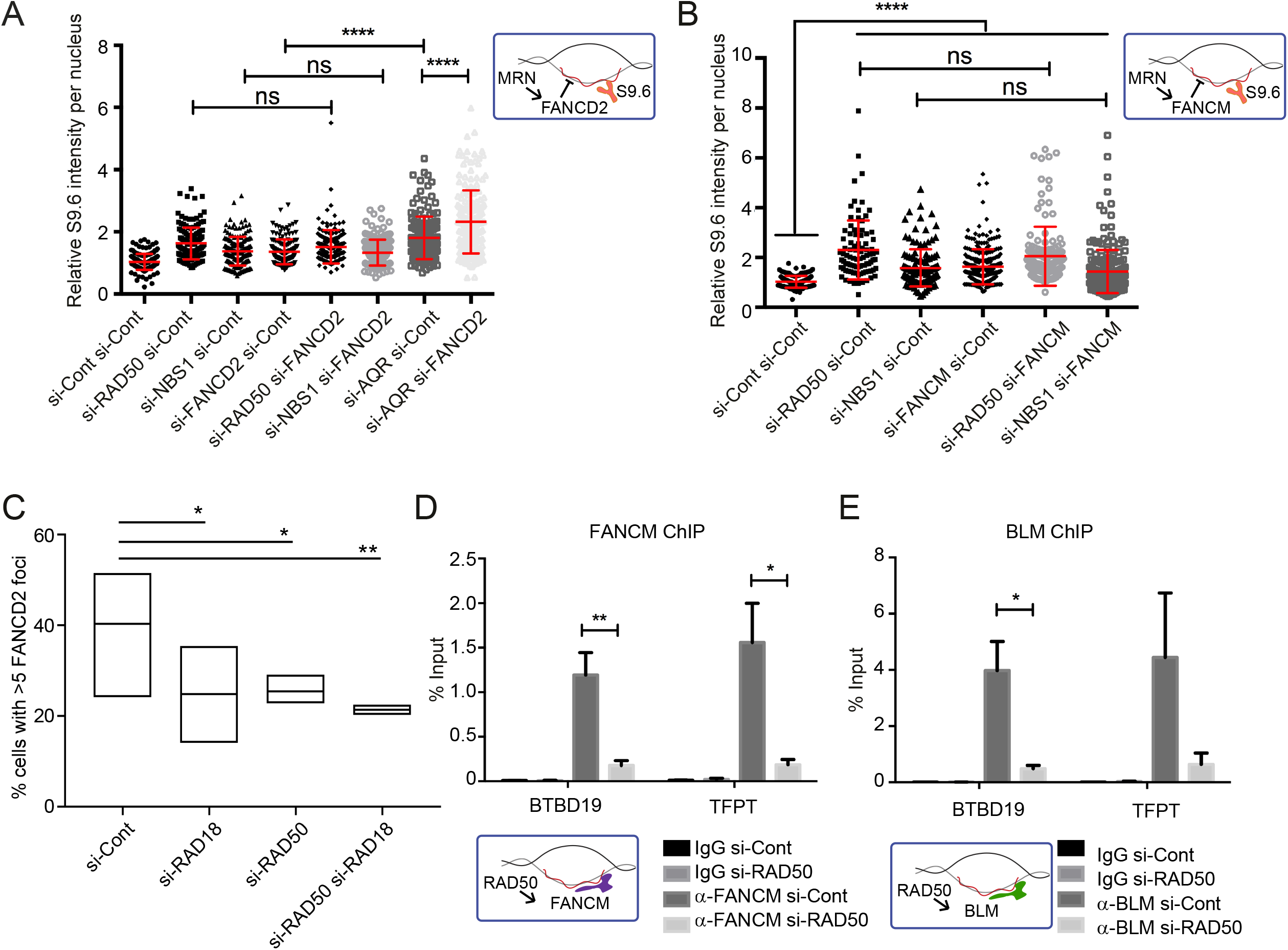
MRN functions in the Fanconi Anemia pathway to recruit FA proteins to R-loops. (A and B) Epistatic effects on nuclear S9.6 staining intensity in the indicated conditions. Aquarius (AQR) is a splicing helicase and R-loop regulator in a different pathway, and AQR depletion showed additive nuclear S9.6 staining increases with si-FANCD2 in (A). N = 3; ANOVA; mean ± SD. (C) Quantification of nuclear FANCD2 foci by immunofluorescence in the indicated cell conditions. Three replicate experiments were performed and the aggregate foci counts were compared by Fisher Exact test and Bonferroni corrected p-value cut-offs were used. (D and E) ChIP of FANCM and BLM at R-loop prone loci BTBD and TFPT depends on RAD50. N = 3; *t* test; mean ± SEM. For all panels, *, P < 0.05; **, P < 0.01; ****, P < 0.0001. Cartoons illustrating the MRN-FA pathway (A,B) and the positive effects of RAD50 on FANCM/BLM recruitment are shown beside each panel.

It has been suggested that the helicase FANCM is a likely direct effector of R-loops and has the ability to unwind R-loops *in vitro* ^13^. To understand how the MRN complex might impact FANCM recruitment to R-loops, we used ChIP-qPCR to show FANCM binding to R-loop prone loci and found that this interaction was abolished when RAD50 is depleted (**Fig. 5D**). This indicates that FANCM recruitment to R-loop sites is significantly less efficient in cells lacking MRN, consistent with the reduction in FANCD2 foci observed above. It has also been suggested that FANCM acts as an anchor and is required for BLM nuclear localization after replication is stalled^44^. Given that BLM is another helicase which we previously demonstrated to impact R-loop accumulation^14^, we performed ChIP-qPCR and also showed that BLM binding to R-loop prone loci requires RAD50 (**Fig. 5E**). These findings support our previous observation that FANCM and BLM have epistatic effects on S9.6 nuclear staining and work in the same R-loop tolerance pathway^14^. Thus, key anti-R-loop helicases are not recruited effectively to R-loop prone sites in MRN-depleted cells.

### MRN loss reduces RAD18 driven PCNA ubiquitination

The observed influence of a non-catalytic role for MRN on R-loop-associated DNA damage and helicase recruitment is novel. However, there are several potential mechanisms linking MRN to recruitment and activation of the FA pathway at R-loop sites. MRN has multiple interaction partners that could directly influence FA activation. One potential route to activation of the FA pathway at replication forks is following PCNA-monoubiquitination (PCNA-Ub) mediated by the E3 ubiquitin ligase RAD18. Previous work showed that PCNA-Ub binds to and stimulates FANCL to monoubiquitinate FANCD2 ^43^. NBS1 has been reported to bind to RAD18 directly ^45^, and yeast MRX components bind to Rad18 and *mre11*Δ deletion strains show reduced PCNA-Ub after genotoxin treatment ^46^. Moreover, *rad18*Δ yeast were found to increase S9.6 staining in our screen (**Fig. 1D**). We first directly tested whether depletion of the human MRN complex subunit RAD50 impacted PCNA ubiquitination. Remarkably, despite increased DNA damage and replication stress upon RAD50 knockdown (**Fig. 2-4**), PCNA-Ub levels on chromatin were reproducibly decreased upon RAD50 depletion (**Fig. 6A**). RAD18, the E3 ligase for PCNA, was depleted as a control and also showed reduced PCNA-Ub on chromatin (**Fig. 6A**). Remarkably, RAD18 foci formation actually increased in RAD50 depleted cells, suggesting that RAD18 does not need RAD50 for localization but that its activity may be impaired in these cells (**Fig. S3A**). Interestingly, RNaseH1 overexpression significantly reduced nuclear RAD18 foci in both control and RAD50 knockdown cells, suggesting that RAD18 may normally localize to R-loop prone sites (**Fig. S3A**). We performed ChIP-qPCR and found that, unlike FANCM and BLM, RAD18 binding to R-loop prone loci does not require RAD50 (**Fig. S3B**).

**Figure 6.**
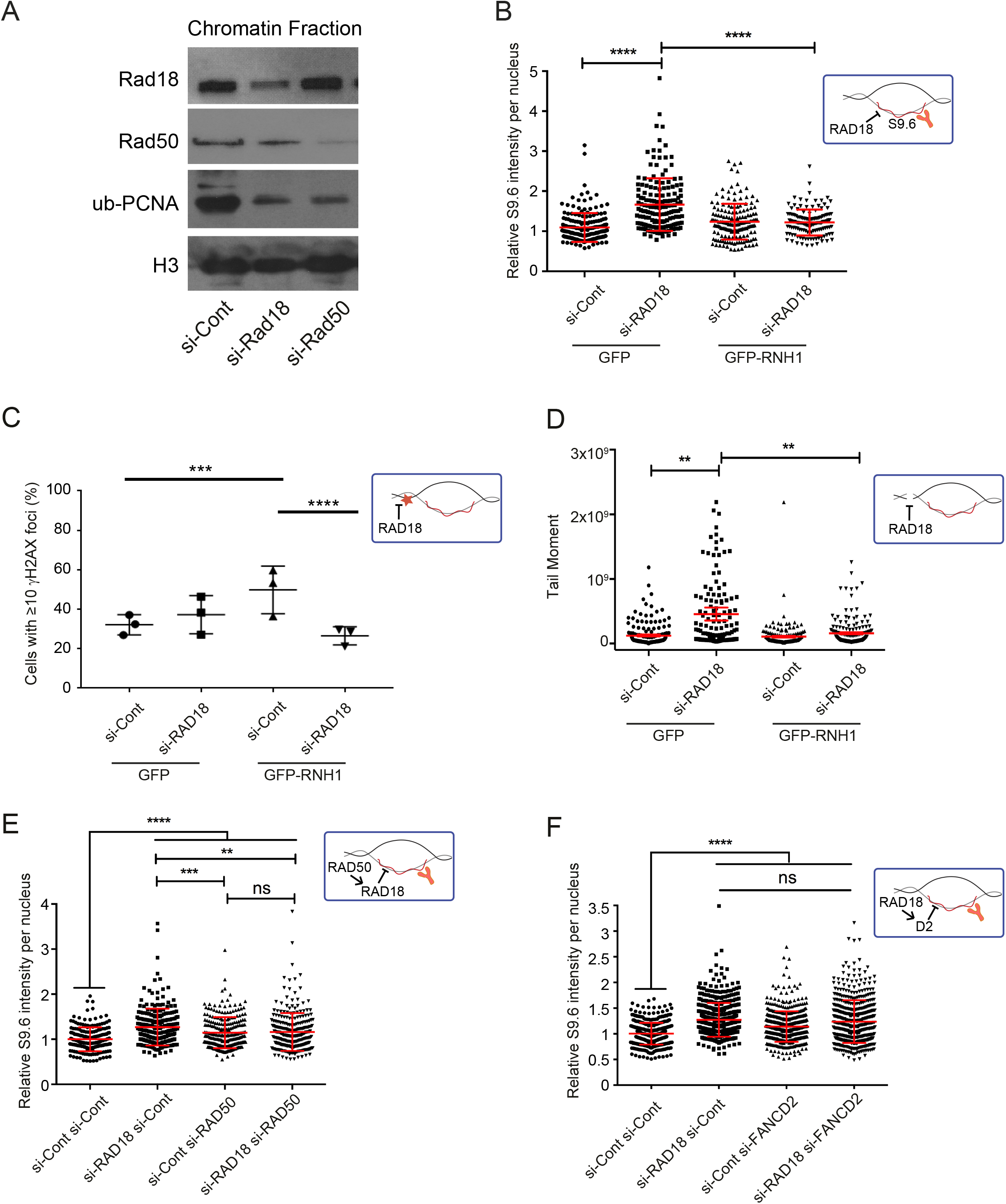
MRN regulates the Fanconi Anemia R-loop tolerance pathway through RAD18. (A) RAD50 depletion inhibited RAD18-dependent PCNA mono-ubiquitination. Representative western blots against the indicated epitopes in the chromatin fraction are shown. Histone H3 was used as loading control for chromatin enrichment. (B) Quantification (top) and representative images (bottom) of S9.6 staining per nucleus in HeLa cells treated with si-Cont or si-RAD18. Cells were transfected with either a control vector (GFP) or one expressing GFP-RNaseH1 (GFP-RNH1). N = 3; *t* test; mean ± SD. (C) RNaseH-dependent γ-H2AX foci induction in RAD18 deficient cells. N = 3; Fisher’s exact test; mean ± SEM. ***, P < 0.001, ****, P < 0.0001. (D) RNaseH-dependent DNA breaks in RAD18 deficient cells. Left, representative comet tail images from single-cell electrophoresis. Right, quantification of comet tail moment under the indicated conditions. N=3; *t* test; mean ± SEM. (E and F) Epistatic effects on nuclear S9.6 staining intensity in the indicated conditions. N = 3 and 4 respectively; ANOVA; mean ± SD. **, P < 0.01; ****, P < 0.0001.

If PCNA modification is an important part of the R-loop tolerance, then RAD18 knockdown should recapitulate R-loop dependent DNA damage phenotypes. Indeed, similar to RAD50 depletion, RAD18 depletion significantly induced S9.6 nuclear staining intensity and DNA damage that was suppressed by co-transfection with GFP-RNaseH1 (**Fig. 6B, 6C** and **6D**). We then performed epistasis experiments via S9.6 nuclear staining of cells doubly depleted for RAD18 and RAD50 or RAD18 and FANCD2 and found that depletion of either combination did not further enhance S9.6 staining intensity comparing to single gene knockdowns (**Fig. 6E** and **6F**, **Fig. S2D** and **S2E**). Finally, depletion of RAD18, like RAD50 and a FANCD2 control, significantly increased RNaseH-sensitive transcription-replication conflicts measured by RNAPII-PCNA proximity ligation (**Fig. S3C**). These data suggest that not only the Fanconi Anemia proteins but also RAD18 act in the same pathway with RAD50 in R-loop regulation. Thus, while MRN is not required for RAD18 localization, it may be needed to stimulate RAD18 to monoubiquitinate PCNA at transcription-replication conflicts to activate the FA pathway and promote the recruitment of anti-R-loop helicases (i.e. FANCM and BLM) and thereby mitigate replication stress (**Fig. 7**, potential models are discussed below).

**Figure 7.**
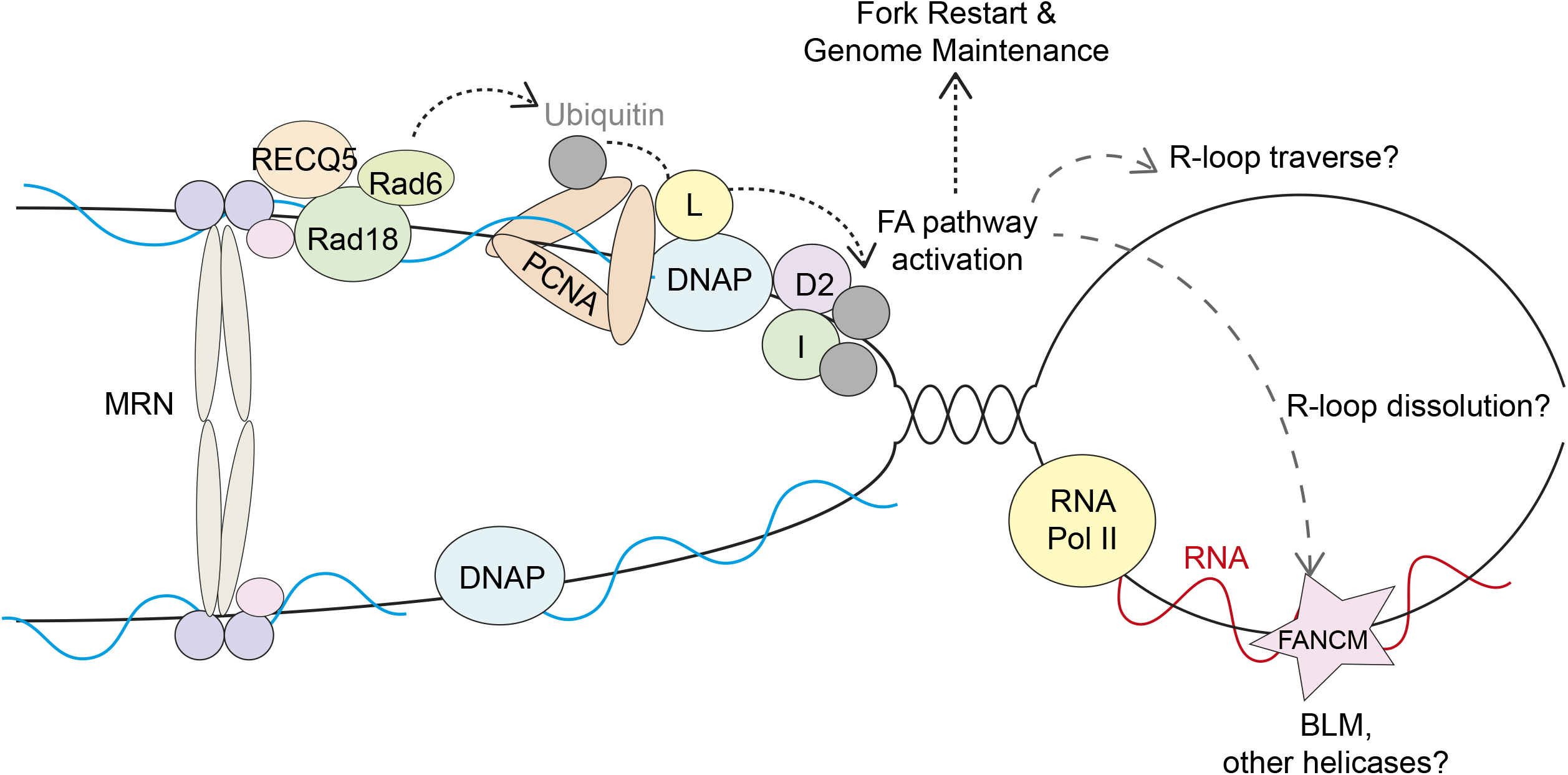
Model of MRN function in R-loop tolerance and mitigation. The MRN complex may act upstream of PCNA ubiquitination and subsequent FA pathway activation and genome stability maintenance. Speculative models for how the FA pathway could resolve these conflicts are indicated with question marks. FANCL, FANCI and FANCD2 are denoted L, I, D2, respectively. See discussion for details.

## DISCUSSION

Ectopic R-loop accumulation can constitute an endogenous source of genotoxic stress that cells must address to mitigate DNA damage and mutagenesis. Here, we use a cross-species approach and identify the MRN complex as an upstream regulator of R-loop-associated DNA damage. We show that recruitment of leading candidates for R-loop removal, FANCM and BLM, to R-loop sites is perturbed in the absence of MRN. The R-loop mitigating activity of MRN is independent of its nuclease function and may rely on its reported SMC-like function in tethering DNA ends ^19^. Furthermore, MRN loss reduces PCNA ubiquitination and is epistatic with knockdown of RAD18, an E3 ubiquitin ligase for PCNA and known activator of the FA pathway^43^. These observations are consistent with the literature in the context of DNA damage repair, where RAD18 requires RECQL5 binding to stimulate PCNA-ubiquitination^47^, and where RECQL5 recruitment to damage sites is dependent on the MRN complex^48^. Our data coalesces with the literature in a model where MRN is an early actor at R-loop stalled forks, and coordinates the recruitment of FA pathway components through a cascade of events (**Fig. 7**).

Over the last few decades, R-loops have been linked to transcriptional pausing ^49^, termination^50^, splicing ^51^, mRNA export ^52^ and other transcription-related phenomena. In this work, we further connect the other component of transcription-replication conflicts, DNA replication, to R-loop tolerance and mitigation. Much of the literature suggests that components functioning in the Fanconi Anemia pathway at replication forks, including BRCA1, BRCA2, BLM, and XPF, are important for R-loop tolerance and removal^2,10,12–14,49,53^. Our data identifies important upstream components to this pathway and raises questions about the unexpected similarities between interstrand crosslink and R-loop bypass or repair. Moreover, other fork-associated proteins implicated in mitigating transcription-replication conflicts, such as RPA, WRN, RECQL5, p53, and MCM2, are not as closely linked to the FA pathway^35,54–57^. This may suggest that there are alternative responses to transcription-replication conflicts that depend on the nature of the R-loop itself or the context of the specific DNA sequence (i.e. replication timing, chromatin state).

In addition, our data suggest for the first time that PCNA modification by RAD18 plays an important role in mitigating R-loop-associated DNA damage. PCNA ubiquitination is critical in the decision to promote translesion synthesis, activate the FA pathway, or promote template switching^43,58^. While MRN has been implicated in PCNA modification by us and others ^46^, the details of this regulation require further study. It is notable that two TLS polymerase subunits, POLD3 and REV3L, have additionally been implicated in preventing R-loop-associated DNA damage^27^. Together, these data suggest that transcription-replication conflicts caused by R-loops are largely handled by the same machinery that deals with chemical DNA damage. What differentiates transcription-replication conflicts from other damage remains unclear and there is much to learn specifically about how pathway choice downstream of PCNA modification is regulated at transcription-replication conflicts.

Finally, it is notable that MRN associates with several helicases that may help to unwind R-loops and facilitate replication fork restart. These helicases include FANCJ, which binds directly to MRN ^59^, WRN which contacts the NBS1 subunit of MRN^38^, RECQL5 which requires MRN to be recruited to DNA damage sites ^48^, and BLM which exhibits interdependence with MRN recruitment depending on the context^37,60^. It remains to be determined how these helicases may cooperate or provide specificity to replisome-R-loop conflicts. Moreover, the role and regulation of other anti-R-loop mechanisms (e.g. Senataxin^61^; XPF^10^) in MRN-deficient cells is unclear. Indeed, it is not known why DNA repair helicases are required when both the replisome and transcriptional apparatus are already associated with multiple helicases capable of unwinding R-loops (e.g. MCM at the replisome, Senataxin, RECQL5, DDX19, DDX21 at RNA polymerase)^50,54,57,62,63^. These activities must collaborate or compete to remove the R-loop and restart the replication fork in a timely manner. Another intriguing possibility is that some of these helicases could facilitate R-loop traverse (**Fig. 7**), where the R-loop is left intact and replication is reinitiated downstream, similar to the role of FANCM and BLM in replisome traverse of interstrand crosslinks ^64^. As we understand more about R-loop tolerance at transcription-replication conflicts, the opportunity to probe the coordination of these events with canonical fork restart mechanisms emerges.

## Acknowledgements

We thank Robert Crouch for the nuclear-targeting GFP-RNaseH1, Iain Cheeseman for the inducible CRISPR/Cas9 RNaseH2A knockout HeLa cells and Hiroyuki Sasanuma for TK6-derived cell lines. P.C.S. is a Canadian Institutes of Health Research (CIHR) New Investigator and a Michael Smith Foundation for Health Research Scholar. J.P.W. holds a CIHR CGS-M scholarship. J.Y.M is a Fonds de la Recherche en Santé du Québec Chair in Genome Stability and was supported by a CIHR Foundation grant. This work was supported by grants from CIHR (MOP136-982), the work was also funded by the Canadian Cancer Society (grant #703263) and a Terry Fox Research Institute New Investigator award (project 1071) to P.C.S.

## Author Contributions

P.C.S and E.Y.C. designed the project. E.Y.C., J.P.W., S.T., Y.A.C, Y.D.Z., and Y.C. conducted the experiments. E.Y.C., P.C.S., J.P.W., S.T., Y.D.Z., Y.A.C., Y.C. and J-Y. M. analyzed the data. L-A. F. and P.H. provided key reagents. P.C.S. and E.Y.C. wrote the manuscript.

## Supplementary Information

**Table S1.** List of SGA results for *rnh1*Δ, *rnh201*Δ and *rnh1*Δ*rnh201*Δ

**Figure S1.** Western blots and comet tail images. (A) Confirmation of RNaseH2A knockout clones by western blot. (B) Western blot showing effective knockdown of the MRN complex in control and dox-induced RNaseH2A deficient HeLa cells. α-tubulin served as loading control. (C) Western blot confirming siRNA knockdown levels of MRN complex from R-loop staining experiments in main text Figure 2B. (D) Representative comet tail images from single-cell electrophoresis in main text Figure 2F.

**Figure S2.** Double siRNA knockdown efficiencies. (A) Western blot showing MRE11 protein levels after 3-day treatment with EtOH (control) or 4-OHT, to induce gene knockout, in the TK6-derived lymphoblasts of the indicated genotype. α-tubulin served as loading control. Related to Figure 4B. (B to E) Western blot confirming double siRNA knockdown levels from R-loop staining experiments in main text Figure 5 A-B and Figure 6 E-F.

**Figure S3.** MRN is not required for RAD18 localization. (A) Depletion of RAD50 increased nuclear RAD18 foci. RNaseH1 overexpression significantly reduced RAD18 nuclear localization in both control and si-RAD50 cells. N = 3; Fisher’s exact test; mean ± SEM. (B) ChIP of RAD18 at R-loop prone loci BTBD and TFPT is independent of RAD50. N = 3; *t* test; mean ± SEM. *, P < 0.05, ****, P < 0.0001.

## Methods

### Yeast genetic interaction screening and validation

Genetic interactions were identified in yeast using synthetic genetic array exactly as described ^22^. A strain was constructed in the SGA query strain Y7092 with *rnh1Δ::NatMX* and *rnh201Δ::URA3* deletions and used for mating and selection of double mutant arrays. Colony size was collected on a flatbed scanner and analyzed using the Balony software suite ^65^. Candidate interactions were recreated by tetrad dissection. Viable double mutants were analyzed by quantitative growth curves using a Tecan M200 plate reader as described ^66^. Gene ontology terms were extracted from the Princeton Generic GO term finder (http://go.princeton.edu/cgi-bin/GOTermFinder). Fold enrichments were calculated by computing the ratio of the number of genes associated with GO term in the hitlist and the number of genes associated with the GO term in the genome. Enriched terms and fold enrichment values were plotted using the web-based ReviGO^67^.

### Yeast chromosome spreads

The indicated mutant strains were grown in rich media and prepared for chromosome spreads and S9.6 staining exactly as described ^25^. Briefly, spreads were probed with a 1:1000 dilution of S9.6 antibody (Kerafast #ENH001), followed by washing in TBS-T and probing with Cy3-labelled anti-mouse antibody. The proportion of nuclei with visible S9.6 staining was scored and the categorical data was tested using a Fisher Exact test then p-values were subjected to a Holm-Bonferroni correction to account for multiple comparisons. To determine the variability in WT, 18 replicates were scored and plotted and the screen was completed in duplicate.

### Cell culture and transfection

HeLa cells were cultivated in Dulbecco’s modified Eagle’s medium (DMEM) (Stemcell technologies) supplemented with 10% fetal bovine serum (Life Technologies) in 5% CO_2_ at 37°C. The control and inducible CRISPR/Cas9 RNaseH2A knockout HeLa cell lines (gifts from Iain Cheeseman lab)^29^ were maintained in DMEM in the absence of doxycycline. To generate stable clones, Cas9 was induced with 1μg/ml doxycycline (Sigma) for 3 days before single-cell sorting. Cells were then expanded in the absence of doxycycline and clones with effective RNaseH2A knockout was screened by Western blotting. The TK6 derived lymphoblasts were gifts from Hiroyuki Sasanuma and were generated and cultured as previously described ^42^. Briefly, TK6 cells were grown in RPMI-1640 medium (Stemcell technologies) supplemented with 5% heat-inactivated horse serum and 200mg/ml sodium pyruvate (Life Technologies). The TK6 derived lymphoblasts were treated with either ethanol (control) or a final concentration of 200nM 4-hydroxytamoxife (4-OHT) (H7904, Sigma) for 3 days in the culture medium to generate the MRE11^−/−^ from MRE11^loxp/loxp^, MRE11^−/WT^ from MRE11^loxp/WT^ and MRE11^−/H129N^ from MRE11^loxp/H129N^.

For RNA interference, cells were transfected with siGENOME-SMARTpool siRNAs from Dharmacon (Nontargeting siRNA Pool #1 as si-Cont, si-RAD50, si-NBS1, si-MRE11, si-FANCD2, si-FANCM, si-AQR, si-RAD18). Transfections were done with Dharmafect1 transfection reagent (Dharmacon) according to manufacturer’s protocol and harvested 48 hours after the siRNA administration. For experiments with overexpression of GFP or nuclear-targeting GFP-RNaseH1 (gift from R. Crouch), transfections were performed with Lipofectamine 3000 (Invitrogen) according to manufacturer’s instructions 24 hours after the siRNA transfections.

### Western blotting

Whole cell lysates were prepared with RIPA buffer containing protease inhibitor (Sigma) and phosphatase inhibitor (Roche Applied Science) cocktail tablets and the protein concentration were determined by Bio-Rad Protein assay (Bio-Rad). Equivalent amounts of protein were resolved by SDS-PAGE and transferred to polyvinylidene fluoride (PVDF) microporous membrane (Millipore), blocked with 5% skim milk in TBS containing 0.1% Tween 20 (TBST), and membranes were probed with the following antibodies: RAD50, γH2AX and FANCM (abcam), NBS1, MRE11, RNaseH2A, RPA2-pS33, RPA2, AQR and RAD18 (Bethyl laboratories), H2AX, p-ATM, p-CHK2, p-ATR and p-Chk1 (cell signaling), FANCD2 (Novus), ub-PCNA(Lys164)(D5C78) (NEB), GFP and GAPDH (Thermo Scientific), and α-tubulin (Life Technologies). Secondary antibodies were conjugated to Horseradish Peroxidase and peroxidase activity was visualized using Chemiluminescent HRP substrate (Thermo Scientific).

### Crystal violet cell viability assay

Control and RNaseH2A CR-knockout cells were counted and seeded in 12-well plates. 48 hours post-siRNA treatment, cells were washed with PBS, fixed with 10% formalin for 30 minutes, and stained with 0.05% crystal violet for 30 minutes at room temperature. The relative levels of stain intensity were quantified by washing the stain with 10% acetic acid in methanol and measuring the absorbance at 590 nm using a TECAN M200. The relative absorbance at 590nm was used to compute expected and observed values based on a multiplicative model of fitness ^68^.

### Immunofluorescence

For S9.6 staining, cells were grown on coverslips overnight before siRNA transfection and plasmid overexpression. 48 hours post-siRNA transfection, cells were washed with PBS, fixed with ice-cold methanol for 10 minutes and permeabilized with ice-cold acetone for 1 minute. After PBS wash, cells were blocked in 3%BSA, 0.1% Tween 20 in 4X saline sodium citrate buffer (SSC) for 1 hour at room temperature. Cells were then incubated with primary antibody S9.6 (1:500) (Kerafast) overnight at 4°C.

Cells were then washed 3 times in PBS and stained with mouse Alexa-Fluoro-568-conjugated secondary antibody (1:1000) (Life Technologies) for 1 hour at room temperature, washed 3 times in PBS, and stained with DAPI for 5 minutes. Cells were imaged on LeicaDMI8 microscope at 100X and ImageJ was used for processing and quantifying nuclear S9.6 intensity in images. For experiments with GFP overexpression, only GFP-positive cells were quantified. For γH2AX and RAD18 foci, the immunostaining was performed the same way except fixation with 4% paraformaldehyde for 15 min and permeabilization with 0.2% Triton X-100 for 5 min on ice for γH2AX (1:1000) (abcam) or fixation with 1% paraformaldehyde for 10 min and permeabilization with 1:1 ration of ice-cold acetone:methanol for 10min for RAD18 (1:500) (Bethyl laboratories). Secondary antibody was rabbit Alexa-Fluoro-568-conjugated antibody (1:1000) (Life Technologies).

### Comet assay

The neutral comet assay was performed using the CometAssay Reagent Kit for Single Cell Gel Electrohoresis Assay (Trevigen) in accordance with the manufacturer’s instructions. Electrophoresis was performed at 4°C and slides were stained with PI and imaged on LeicaDMI8 microscope at 20X. Comet tail moments were obtained using an ImageJ plugin as previously described ^69^. At least 50 cells per sample were analyzed from each independent experiment.

### Native BrdU

Cells were grown on coverslips overnight before siRNA transfection and plasmid overexpression. 48 hours post-siRNA transfection, a final concentration of 30 μM BrdU (Sigma) was added to the culture medium and incubated for 24 hours at 37°C. After a 24 hour incubation with BrdU, cells were rinse with PBS twice, fixed with ice-cold 70% ethanol for 10 minutes and blocked with 3% BSA, 0.1% Tween 20 in 4X SSC for 1 hour at room temperature. Cells were than incubated with anti-BrdU antibody (1:500) (BD) for 1 hour at room temperature, washed 3 times in PBS and stained with mouse Alexa-Fluoro-568-conjugated secondary antibody (1:1000) (Life Technologies) for 1 hour at room temperature. Cells were then stained with DAPI for 5 minutes and imaged on LeicaDMI8 microscope at 100X. ImageJ was used for processing and quantifying nuclear BrdU intensity in images.

### Proximity Ligation Assay

Cells were grown on coverslips, washed with PBS, fixed with 4% paraformaldehyde for 15 minutes and permeabilized with 0.2% Triton X-100 for 5 minutes. Cells were then blocked in 3% BSA, 0.1% Tween 20 in 4XSSC for 1 hour at room temperature. Cells were then incubated with primary antibody overnight at 4°C [1:500 goat anti-RNA polymerase II antibody (PLA0292, Sigma) with 1:500 rabbit anti-PCNA antibody (PLA0079, Sigma)]. The next day after washing with 1XPBS twice, cells were incubated with pre-mixed PLA probe anti-goat minus and PLA probe anti-rabbit plus (Sigma) for 1 hour at 37°C. The subsequent steps in proximal ligation assay were carried out with Duolink In Situ Kit (Sigma) in accordance to manufacturer’s instructions. Slides were then stained with DAPI and imaged on LeicaDM18 microscope at 100X. Negative controls were treated identically but anti-RNA polymerase II antibody was omitted.

### Genomic DNA extraction and DNA combing

Cells were first pulse labeled for 20 minutes with 250μM IdU (Sigma), washed twice with PBS, and then pulse labeled with 30μM CldU (Sigma) for 20min. Cells were then collected and scraped into ice-cold PBS and genomic DNA was extracted with CombHeliX DNA Extraction kit (Genomic vision) in accordance with the manufacturer’s instructions. DNA fibers were stretched on vinyl silane-treated glass coverslips (CombiCoverslips) (Genomic vision) with automated Molecular Combing System (Genomic Vision). After Combing, the stretched DNA fibers were dehydrated in 37°C for 2 hours, fixed with MeOH:Acetic acid (3:1) for 10 minutes, denatured with 2.5 M HCl for 1 hour, and blocked with 5% BSA in PBST for 30 minutes. IdU and CldU were then detected with the following primary antibodies in blocking solution for 1 hour at room temperature: mouse anti-BrdU (B44) (1:40) (BD) for IdU and rat anti-BrdU [Bu1/75 (ICR1)] (1:50) (abcam) for CldU. After PBS wash, fibers were than incubated with secondary antibodies anti-Rat-Alexa488 (1:50) (Invitrogen) and anti-mouse-Alexa 568 (1:50) (Life Technologies) for 1 hour at room temperature. DNA fibers were analyzed on LeicaDM18 microscope at 100X and ImageJ was used to measure fiber length. For the experiment with transcription inhibitor, cells were first treated with DMSO or 50μM cordycepin for 2 hours before IdU labeling.

### FACS analysis for S9.6

Cells were harvested, washed with 1XPBS twice, extracted with 25mM HEPES, 50mM NaCl, 1mM EDTA with protease inhibitor on ice, and fixed with 2% paraformaldehyde in 1x PBS ^70^. Cells were then incubated with primary antibody S9.6 (1:200) (millipore) overnight at 4°C and with anti-mouse-IgG-PerCP-Cy5.5 secondary antibody (1:100) (Santa Cruz) for 30 minutes at room temperature. Cells were resuspended in 1XPBS and analyzed with a FACSCalibur flow cytometer using CellQuestPro software. The mean S9.6 intensity and histograms were generated using FlowJo Version 9.3.2.

### DRIP and ChIP-qPCR

The method was followed from ^71^ with some modifications. Cells were crosslinked in 1% formaldehyde for 10 minutes before quenching with glycine for 5 minutes at room temperature, and then lysed in ChIP lysis buffer (50mM HEPES-KOH at pH 7.5, 140 mM NaCl, 1 mM EDTA at pH 8, 1% Triton X-100, 0.1% Na-Deoxycholate, 1% SDS) and rotated for 1 hour at 4 °C. DNA were sonicated on Q Sonica Sonicator Q700 for 8 minutes (30 sec ON, 30 sec OFF) to generate fragments of 200-500bp. For DRIP, the chromatin preps were treated with 20 mg/mL Proteinase K (Thermo Fisher Scientific) at 65°C overnight and total DNA was purified by phenol/chloroform purification method. Protein A magnetic beads (Bio-rad) were first pre-blocked with PBS/EDTA containing 0.5% BSA and then incubated with S9.6 antibody in IP buffer (50 mM Hepes/KOH at pH 7.5; 0.14 M NaCl; 5 mM EDTA; 1% Triton X-100; 0.1% Na-Deoxycholate, ddH2O) at 4°C for 4 hour with rotation. DNA was then added to the mixture and gently rotated at 4°C overnight. Beads were recovered and washed successively with low salt buffer (50mMHepes/KOH pH 7.5, 0.14 M NaCl, 5 mM EDTA pH 8, 1% Triton X-100, 0.1% Na-Deoxycholate), high salt buffer (50 mM Hepes/KOH pH 7.5, 0.5 M NaCl, 5 mM EDTA pH 8, 1% Triton X-100, 0.1% Na-Deoxycholate), wash buffer (10 mM Tris-HCl pH 8, 0.25 M LiCl, 0.5% NP-40, 0.5% Na-Deoxycholate, 1 mM EDTA pH 8), and TE buffer (100 mM Tris-HCl pH 8, 10 mM EDTA pH 8) at 4°C, two times. Elution was performed with elution buffer (50mMTris-HCl pH 8, 10mMEDTA, 1% SDS) for 15 minutes at 65°C. After purification with PCR Cleanup kit (Sigma-Aldrich), nucleic acids were eluted in 100 μL of elution buffer (5 mM Tris-HCl pH 8.5) and analyzed by quantitative real-time PCR (qPCR). For ChIP, DNA and antibody were incubated in IP buffer overnight with rolling at 4°C. Antibody-bound DNA was recovered using the Protein A or Protein G magnetic beads (Bio-Rad), washed similarly as DRIP samples and treated with Proteinase K and RNAse after elution. Then, antibody-bound DNA was purified with PCR Cleanup kit and analyzed by qPCR. qPCR was performed with Fast SYBR Green Master (ABI) on AB Step One Plus real-time PCR machine (Applied Biosystem). Primer sequences are BTBD19-Forward (5’-CCCCAAAGGGTGGTGACTT), BTBD19-Reverse (5’-TTCACATTACCCAGACCAGACTGT), TFPT-Forward (5’-TCTGGGAGTCCAAGCAGACT) and TFPT-Reverse (5’-AAGGAGCCACTGAAGGGTTT). qPCR results were analyzed using the comparative CT method. The RNA-DNA hybrid and ChIP DNA enrichments were calculated based on the IP/Input ratio.

### Cell fractionation

Cells were washed with cold PBS, resuspended with buffer A (10mM Hepes-NaOH, pH 7.9, 10mM KCl, 1.5mM MgCl2, 0.34 M sucrose, 10% glycerol, and 1mM DTT) with Triton X-100 and incubated on ice for 5 minutes. Nuclei were collected by centrifugation at 1500g for 4 minutes at 4°C, washed with buffer A with Triton X-100 twice, resuspended in buffer B (3mM EDTA, 0.2 mM EGTA and 1mM DTT) and incubated on ice for 15 minutes. Samples were then centrifuged at 13000rpm for 15 minutes at 4°C. The supernatant (nucleoplasmic fraction) were separated from the pellet (chromatin fraction), and the pellet was resuspended in buffer B and sonicated for 10 minutes in cold water.

### MRN nuclease assays

The substrates used were DNA/DNA hybrid (JYM3395 GTTTCTGGACCATATGATACATGCTCTGGCCAAGCATTCCGGCTGGTCGCTAATCGTTGA and JYM3393 TCAACGATTAGCGACCAGCCGGAATGCTTGGCCAGAGCATGTATCATATGGTCCAGAAAC) or DNA/RNA hybrid (JYM3395 GTTTCTGGACCATATGATACATGCTCTGGCCAAGCATTCCGGCTGGTCGCTAATCGTTGA and JYM3394 rUrCrArArCrGrArUrUrArGrCrGrArCrCrArGrCrCrGrGrArArUrGrCrUrUrGrGrCrCrArGrArGrCrArUrGrUrArUrC rArUrArUrGrGrUrCrCrArGrArArArC). JYM3395 was end-labelled in 5’ with γ-ATP and T4 polynucleotide kinase (NEB) according to the manufacturer’s conditions. The complementary strand was annealed and the DNA/DNA or DNA/RNA hybrids were purified from an 8% PAGE gel.

MRN exonuclease reactions were performed in 25 mM MOPS (morpholinepropanesulfonic acid; pH 7.0), 60 mM KCl, 0.2% Tween-20, 2 mM dithiothreitol, 2 mM ATP and 5 mM MnCl_2_, with 100 nM DNA/DNA hybrid or DNA/RNA hybrid radiolabeled substrates. Reaction mixtures were incubated for 1 hour at 37°C, followed by treatment with one-fifth volume of stop buffer (20 mM Tris–Cl pH 7.5 and 2 mg/mL proteinase K) for 30 min at 37°C. Formamide loading buffer was added to the reaction mixtures, which were heated at 95°C for 5 min and then loaded onto 10% polyacrylamide denaturing gels. Gels were migrated at 55 W for 120 min and then exposed in a phosphorimager cassette or on film. The percentages of DNA resection, from three independent experiments, was calculated from the intact DNA substrate in the reaction mixtures divided by the intact DNA substrate in the no enzyme control. This provided the percentage of DNA remaining after resection which was substracted from 100.

